# Meditation in the third-person perspective modulates minimal self and heartbeat-evoked potentials

**DOI:** 10.1101/2024.10.24.619995

**Authors:** Hang Yang, Bruno Herbelin, Chuong Ngo, Loup Vuarnesson, Olaf Blanke

## Abstract

Experienced meditation practitioners often report altered states of their sense of self, including decentering and distancing the self from the body and one’s current concerns. Altered states of the sense of self, such as disembodiment and distancing of the self from the body, have also been induced experimentally using virtual reality (VR) and linked neurally to heartbeat evoked potentials (HEPs). Whereas many studies investigated the related neural correlates of such decentering during meditation, none experimentally modulated the sense of self during meditation practice using VR nor determined the potentially associated behavioral changes of the sense of self. Here we determined HEPs and behavioral measures in 23 participants who performed a guided meditation practice in VR, either from a third-person (3PP) or first-person perspective (1PP) to modulate the sense of self. In the 3PP versus 1PP meditation condition, we report stronger sensations of detachment and disconnection, reduced salience of the perceived body boundary, and reduced self-identification with the body. HEP analysis revealed differential neural responses between conditions, characterized by a more negative HEP amplitude in the 3PP condition, associated with activation of the posterior cingulate cortex and medial prefrontal cortex. Leveraging a new VR-supported meditation platform and methods, these data link the sense of self in meditation practice to the neuroscience of the bodily self, based on subjective, behavioral, and neural measures.

## 1. Introduction

Inquiring about the nature of the self is a fundamental part of many contemplative traditions (i.e., Albahari, 2006; Dahl et al., 2015; Hölzel et al., 2011; Kessel et al., 2016; Siderits et al., 2011; Tang et al., 2015). Heightened focus on the self in daily life (i.e., self-related thoughts, autobiographical memories, personal goals, mental time travel), has been criticized by these traditions and argued to lead to psychological distress (Dahl et al., 2015; Nikitin & Freund, 2021; Vago & Silbersweig, 2012) and meditation practitioners are often guided to distance and detach themselves from their conscious experience and self-related cognitive activities. Several contemplative practices encourage practitioners to take a distanced and de-centered perspective and to remain as a non-judgmental observer of ongoing fluctuating conscious experience (Dahl et al., 2015; Hölzel et al., 2011; Kessel et al., 2016; Millière et al., 2018), which often concern the practicioners’s body and/or breath(Anālayo, 2019; Brandmeyer & Delorme, 2018; Tang et al., 2015).

Research in cognitive neuroscience also investigated perspective taking and resonates with this dichotomy between higher-level self-rated cognitions versus lower-level bodily perceptions, distinguishing two main forms of self-consciousness: a narrative self and a minimal (bodily) self (Damasio, 1999; Gallagher, 2000; Legrand & Ruby, 2009; Millière et al., 2018; Northoff et al., 2006; Seth & Tsakiris, 2018). The narrative self refers to a higher-level cognitive self, involving self-related thoughts, language and autobiographical memories, that shape our life narrative and core self-related beliefs. The minimal self, instead, is not extended in time and centers on conscious bodily experiences such as body ownership and self-identification with the body, based on multisensory perception. The minimal self can be altered experimentally leading to changes in body ownership and self-identification through the manipulation of specific multisensory bodily signals(Blanke et al., 2015; Park & Blanke, 2019). In such multisensory paradigms using virtual reality (VR) (Bayer et al., 2023; Ehrsson, 2007; Lenggenhager et al., 2007; Noel et al., 2015; Salomon et al., 2016), participants receive touches on their backs while viewing an avatar from a third person perspective (3PP; seen from behind and projected in front of them and receiving the same tactile stimulation), inducing illusory self-identification with the avatar.

Whereas, the narrative self and related processes have been studied in meditation(i.e., Britton et al., 2021; Farb et al., 2007; Vago & Zeidan, 2016), only a few researchers studied the role of the minimal self in meditation and how it depends on multisensory bodily mechanisms and a de-centered 3PP. Based on in-depth interviews of long-term meditators, it was reported that mindfulness meditation modulates body ownership and the perceived body boundary (Ataria et al., 2015; Dor-Ziderman et al., 2016). These changes were associated with beta oscillations in a large cortical network, centered over right temporo-parietal cortex (Dor-Ziderman et al., 2013, 2016), a region known to be involved in self-identification and a shift from the first-person perspective (1PP) to a 3PP (Blanke et al., 2002, 2005; Ionta et al., 2011). Related more recent work also reported changes in self-location (Dambrun et al., 2023) and 1PP in meditation practice (Nave et al., 2021) and highlighted beta-band oscillatory changes in the medial parietal cortex (Trautwein et al., 2024). While these studies launched important research on specific aspects of the minimal self in meditation, they did neither experimentally manipulate nor behaviorally measure changes of the minimal self (such as changes in self-identification with the body) during the recorded meditation sessions, relying instead on in-depth verbal interviews about subjective experiences that are, however, prone to participant and experimenter biases (i.e., Adler, 1973; Rosenthal & Fode, 1963).

Here, we developed a meditation research platform using immersive virtual reality (VR) allowing us to experimentally manipulate the minimal self during ongoing meditation practice (3PP versus 1PP meditation conditions) and to measure changes of the minimal self (self-identification, perceived body boundary) immediately after the meditation session. In the 3PP versus 1PP meditation condition, we report stronger sensations of detachment and disconnection, reduced salience of the perceived body boundary, and reduced self-identification with the body. Heart-beat evoked potentials (HEPs) recorded during meditation revealed a more negative HEP amplitude during 3PP meditation.

## 2. Materials and Methods

### 2.1. Participants

We tested a group of 23 right-handed healthy participants (12 female, 11 male) for the study. Sample size was determined based on the effect size (0.72) for the HEP effect between synchronous and asynchronous conditions from Park et al. (2016). A paired t-test with an alpha level of 0.05 and a power of 0.9 was assumed for the estimation of the sample size (G*Power 3.1.9.7 software, Faul et al., 2007). Participants were all right-handed and proficient in English, aged from 19 to 48 years (mean: 28.7 years, standard deviation: 7.6 years). They were all non-meditators who reported no prior experience with meditation practice. Exclusion criteria were the presence of neurological or psychiatric disorders and substance abuse. Participants gave informed consent before the experiment, and a compensation of 20 CHF per hour was paid after completion of the experiment. The study was approved by the Swiss Ethics Committees on research involving humans (Project ID: 2021-00358).

### 2.2. Virtual-reality experimental setup

The VR immersion platform developed for this study includes an in-house real-time volumetric body scanning system (Albert et al., 2019) using 4 cameras (Kinect Azure cameras, Microsoft, Washington, USA) mounted on a metallic frame, a VR headset for visual immersion (HP Reverb G2, Hewlett-Packard, USA) with 107° diagonal FOV and a resolution of 2160×2160 pixels per-eye), and earplugs for audio immersion (DH Gold series, Italy). The experiment was developed with our custom-made open-source software ExVR (https://github.com/lnco-epfl/ExVR) which uses Unity3D (Unity Technologies, USA) for real-time 3D rendering.

Two audio-visual virtual environments were designed for the experiment. The virtual laboratory environment (Figure 1A) was a 3D replica of the experimentation room; it was used for welcoming participants into VR and as an intermediate place where they answered questionnaires and performed tasks while still immersed. The virtual forest environment represented an outdoor scene for immersing participants in a natural landscape for the meditation (i.e., trees and vegetation moving with the wind; birds chirping; sound of river flowing in the background based on custom binaural audio recordings in the mountains).

**Figure 1.**
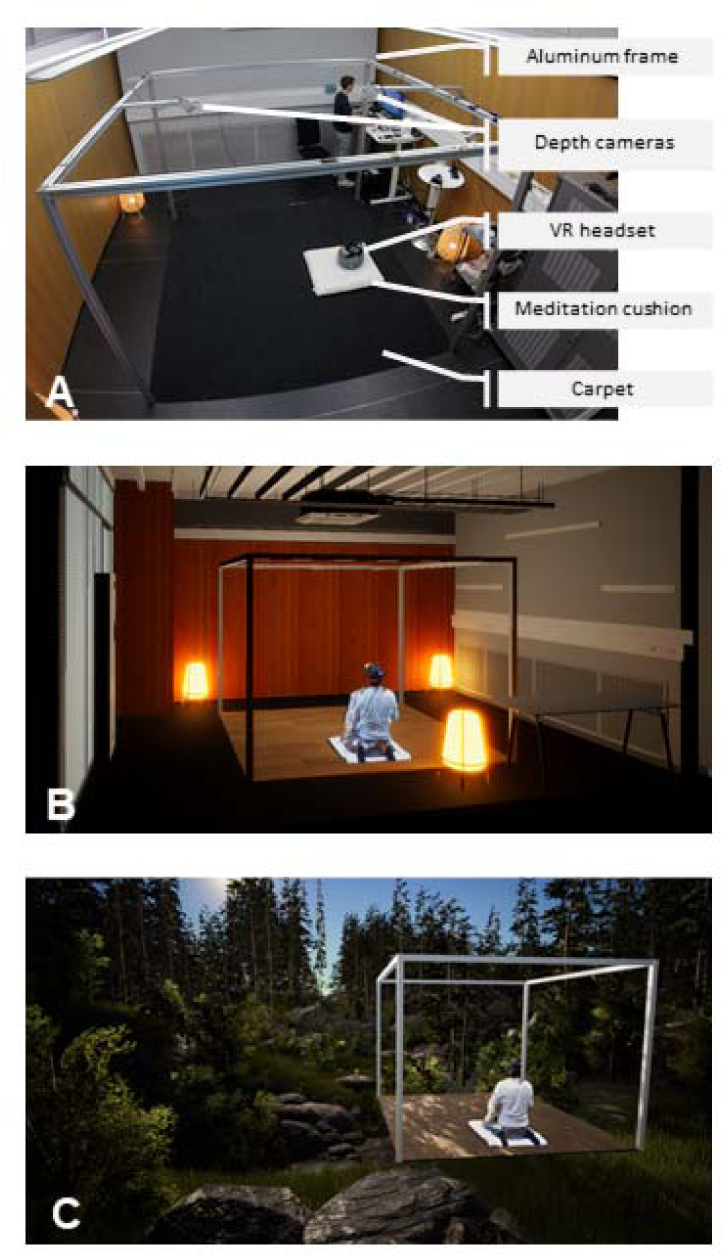
Experimental setup and mixed-reality environment for the immersive meditation experiments. A) a virtual laboratory room and equipment, B) a meditator sitting in the virtual laboratory room C) a meditator sitting in the virtual forest for meditation.

Our body scanning technology allows reconstructing a 3D representation of the body of participants located in the field of view of the cameras, either in real-time or by replaying pre-recorded 3D captures(Albert et al., 2019; Vuarnesson et al., 2024). The VR platform thereby allowed immersing participants in different virtual environments where they could see a 3D representation of their own body (collocated with their physical body) while the movements of their body were captured and shown in real-time with their actual movements (Figure 1B, 1C; see Vuarnesson et al., 2024) for video examples). 3D representations of other people (such as a person providing the meditation instructions) could also be integrated into the virtual environments (Figure 1C) and this person could be physically co-present with participants during the experiment (i.e. real-time capture) or be a previously recorded person.

### 2.3. Experimental design and procedure

The experiment followed a single factor within-subject design to compare subjective reports, behavioral and physiological measurements between two conditions involving a 15-minute meditation session in immersive VR. The order of the two meditation conditions (3PP, 1PP) was randomized across participants. Three short questionnaires and one behavioral task were performed inside VR immediately after both meditation conditions for evaluating differences in disembodiment, mindfulness, and perceived body boundary (see below for details). A more extensive questionnaire was administered after each meditation session, and outside of VR, to assess our participants’ states of consciousness during meditation more globally. The physiological signals recorded during the whole experiment included electroencephalogram (EEG), electrocardiogram (ECG), and respiration.

The experiment took place in a dedicated room furnished to provide a relaxing atmosphere; dimmed lights, a square carpet on which participants walked without shoes, and soft cushions at the center on which they were sitting during the experiment. Once comfortably installed, participants were equipped with the VR headset and earplugs. Participants were first immersed in the virtual laboratory environment (a virtual room looking like the actual laboratory room) and habituated to the interaction modalities and to the view of their own avatar. After a few minutes of habituation they were asked to fill the Perceived Body Boundary Questionnaire (PBBQ), which reflects the perceived salience of the body boundary and has been used before in meditation research (Dambrun, 2016; Hanley et al., 2020).

Participants then ‘met’ with a virtual meditation guide (a professional meditation teacher prerecorded in the same virtual room using the same VR system) inviting them to carry out the meditation session in a different VR ‘place’. After a short transition period (2 seconds), participants found themselves immersed in the virtual forest environment (Figure 2A). They were then instructed to follow the verbal guidance, provided by the prerecorded virtual meditation guide. After an invitation to visually explore the virtual forest environment, they were asked to close their eyes and were then guided through a mindfulness exercise designed to help them calm their mind and focus their awareness on their body. The 3PP condition consisted of focused awareness on different parts of the participant’s body, with particular emphasis on the torso (5 minutes). After this, participants were guided through an open monitoring meditation(i.e., Hölzel et al., 2011; Kessel et al., 2016; Lutz et al., 2008) during which they were instructed to take a de-centered and distanced perspective and witness moment-to-moment fluctuations occurring in their conscious experience. Up to this point, participants had always experienced the virtual forest and their body from their habitual embodied viewpoint. Now this viewpoint was slowly and progressively shifted to a third-person perspective, by placing their virtual viewpoint to a slightly elevated and posterior position behind their virtual body (Figure 2B). After an instruction to witness their body from this 3PP, the guidance continued with open monitoring. The 1PP condition had the same length but differed in two main aspects from the 3PP condition. Following the focused awareness on different body parts, (1) participants followed meditation guidance that centered on external objects in the forest (i.e., attention was focused on objects in the virtual forest such as trees and rocks, and not on body and self as in the 3PP meditation condition). In addition, (2) there was no shift in viewpoint during meditation and participants remained in 1PP. We controlled visually for the visual displacement (of the viewpoint during 3PP) by moving during 1PP the entire shelter area within the virtual forest and by displacing it by the same amplitude/distance and with the same speed as the 3PP condition (but now participants remained in 1PP) (see Fig. 2).

**Figure 2.**
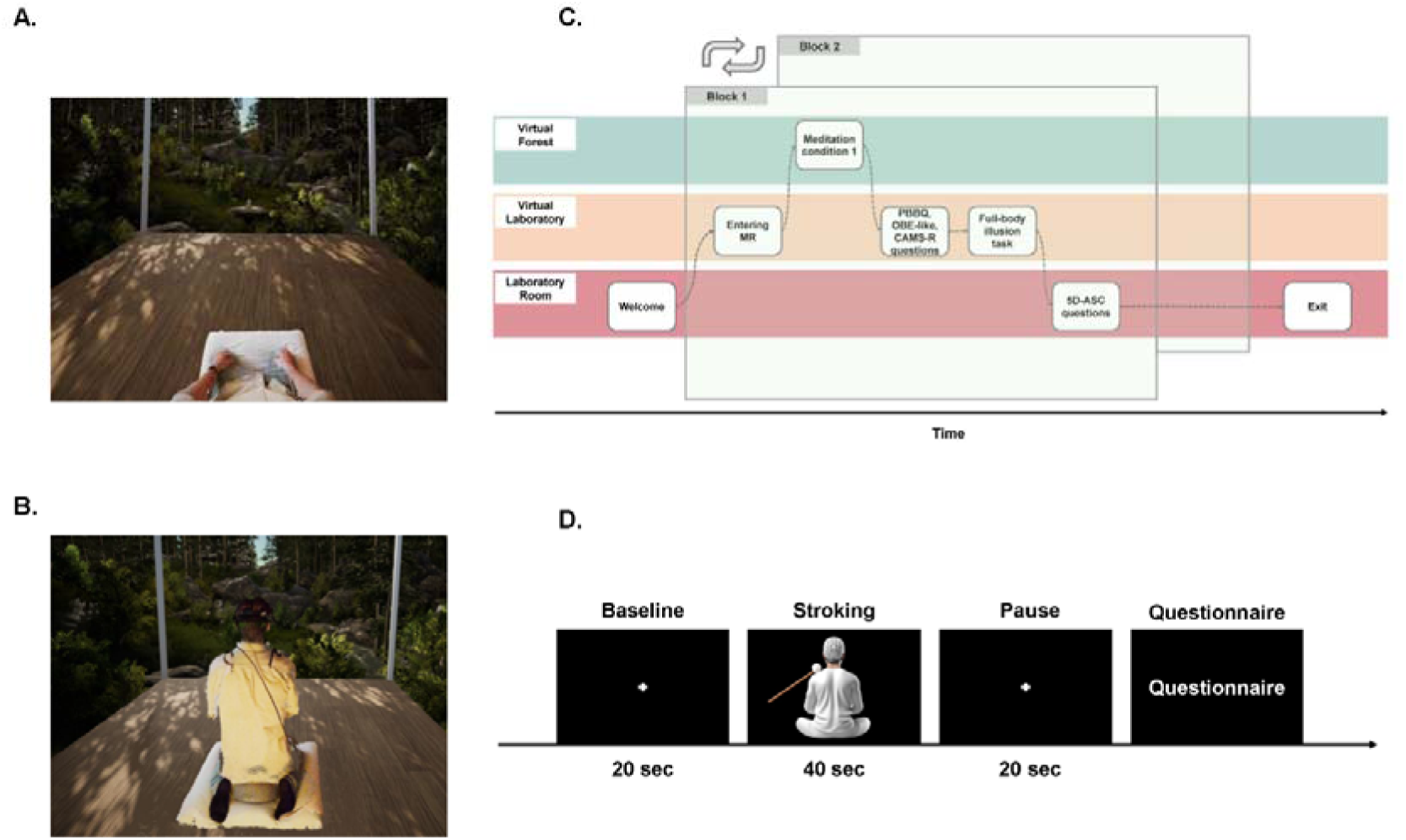
A) Participants see their bodies in first-person view when immersed in the virtual forest environment. B) In the 3PP condition, the visual perspective is manipulated to show the participant’s body in a third-person view. C) A simplified experiment flow of one typical block in the study. The experiment consists of two blocks, one per condition (3PP or 1PP). In each bloc, the meditation sessions were followed by subjective evaluations with questionnaires in MR (PBBQ, OBE, and CAMS-R), a behavioral task in MR (full body illusion), and a last questionnaire outside MR (5D-ASC). The order of the two meditations was counterbalanced across participants. D) Illustration of one block of the FBI task.

After each meditation condition, participants first returned to the virtual laboratory where they were asked to answer several questions while still immersed in VR. The PBBQ was asked again so as to compare its score with the baseline measured before meditation. This was followed by the Cognitive and Affective Mindfulness Scale (CAMS-R), to captures mindfulness states that reflects well-being and emotion regulation as a dynamic state (Feldman et al., 2007, 2022), and by an Out-of-Body Experience sensation questionnaire (OBE), developed in-house to evaluate subjective aspects of out-of-body experiences (Song, 2020; Wu et al., 2023). For statistical analysis of the questionnaire responses, we used a Linear Mixed Effect (LME) model (significance level p<0.05, two-tailed) and Bayes Factor (implemented by the BayesFactor package, Morey et al., 2011) to compare the difference between 3PP condition and 1PP condition.

Next, a full-body illusion task was then administered to evaluate potential differences in sensitivity to multisensory visuo-tactile stimulation between the 3PP and 1PP conditions. During the full-body illusion task, participants see a representation of their body as seen from a position behind their body while their back is stroked by the experimenter (Aspell et al., 2009; Bayer et al., 2023; Lenggenhager et al., 2007, 2009; Park et al., 2016). The present full-body illusion was adapted to the sitting position of the participants in the VR environment and the stroking was applied with a stick ending with a white ball held by the experimenter (unaware of the condition). We tested the full-body illusion in two conditions, in a synchronous visuo-tactile stimulation (i.e. participants saw the ball touching their back synchronously with the moment when they felt the touch on their back) and in an asynchronous condition (i.e. participants saw the ball touching their back with a 500 ms delay to the moment when they felt the touch on their back). The synchronous and asynchronous conditions were each repeated twice in randomized order. All four full-body illusion trials started by displaying a fixation cross in front of participants for 20 seconds, followed by 40 seconds of visuo-tactile stimulation (see Figure 2D). At the end of each trial (as in Aspell et al., 2009; Park et al., 2016), we gauged participants’ mislocation of touch ( “I felt as if the touch was located where I saw the stroking”), self-identification ( “I felt as if the body I saw was my body”), and a control question ( “I felt as if I had more than three bodies”), using a 7-point Likert scale.

Finally, participants took off the VR headset and earplugs and were given a tablet on which to fill the 5-Dimensional Altered States of Consciousness Rating Scale questionnaire (5D-ASC, Schmidt & Berkemeyer, 2018). After a short break, the other condition was performed following a similar procedure.

### 2.4. EEG recording

For the entire duration of the experiment, EEG was recorded using a 64-channel eego^*TM*^ mylab system (ANT Neuro). Based on the five percent electrode placement system (Oostenveld & Praamstra, 2001), a total of 65 EEG channels were placed. These included two mastoid electrodes placed behind the ears (M1 and M2), the CPz electrode selected by the system as the online reference of all channels, and the AFz electrode used as the ground electrode. The impedance of all EEG channels was kept below 20 kΩ for the whole experiment. The Lab Recorder was used for online recordings of the Lab Streaming Layer (LSL, https://github.com/sccn/labstreaminglayer) at a sampling rate of 512 Hz.

### 2.5. ECG and respiration recordings

ECG and breathing signals were simultaneously recorded through bipolar channels connecting to the EEG system. ECG signals were collected with two bipolar ECG channels diagonally placed, one attached between the lower two ribs on the left side of the body, and the other under the right collarbone around 5 cm from the sternum. Breathing was recorded with a respiration belt (Spec Medica) attached to the lower chest (thoracic diaphragm region) of the participants.

### 2.6. Analysis of the heartbeat evoked potentials

All analyses of ECG, EEG, and respiration signals were done in MATLAB (The MathWorks, Inc.). EEG signals were analyzed using EEGLAB (Delorme & Makeig, 2004) and fieldtrip (Oostenveld et al., 2011).

#### ECG pre-processing

The ECG signal was first filtered with high-pass and low-pass frequency (0.1 – 40 Hz). To detect the R-peak of heartbeats, the filtered ECG signals were decomposed into 5 scales based on the maximal overlap discrete wavelet transform (Biran & Jeremic, 2023). The ECG waveforms were reconstructed with wavelet coefficients of only the fourth and fifth scale which corresponded to 11.25 - 22.5 Hz and 5.625 – 11.25 Hz. The R-peak detection was applied to find the peak time of the squared values of the reconstructed signals. The peak detection results were visually inspected throughout the experiment for all participants. A threshold of 10000 microvolts in ECG amplitude lasting for more than 200 s was set for all participants to exclude artifacts caused by motion or pull of ECG cables during the experiment. Detected noisy periods were excluded from further analysis.

#### EEG pre-processing

Bad channels were marked due to an unstable EEG signal during the setup of the EEG cap, only one channel for one participant was removed and was interpolated with the average amplitudes of surrounding channels. Continuous EEG was filtered with the same bandpass filter as for ECG (0.1 – 40 Hz), followed by segmenting the continuous EEG into epochs from 100ms before to 800ms after the R-peak. EEG epochs were baseline-corrected with the pre-stimulus time [-100, 0] *ms*, as widely used for HEPs studies (Coll et al., 2021; Kumral et al., 2022; Terhaar et al., 2012). Trials with absolute values in their amplitudes higher than 100 microvolts were automatically detected and removed for further analysis. This resulted in an average of 787 (SD: 156) trials during the 3PP condition and an average of 730 (SD: 146) trials for the 1PP condition. The remaining EEG trials were then re-referenced to the average of the mastoid channels, as in previous HEPs studies (Coll et al., 2021). Importantly, the mastoid channels were chosen as the re-reference (instead of the common average), in order to diminish the impact of alpha-band oscillations in the posterior sites (which was observed in our data such as electrodes O1, PO7, etc.), as has been supported by numerous meditation studies (Katyal & Goldin, 2021; Lee et al., 2018; Lomas et al., 2015). The EEG trials were then fitted into an independent component analysis (ICA) and the waveform and topography of all components were visually inspected. The components related to eye blinks and saccades were removed. An average of 1.48 (*SD*: 0.73) independent components were removed in the ICA. The clean EEG trials were then averaged separately for both 1PP and 3PP conditions for each participant.

#### HEPs statistical analysis

To test whether and when the HEPs showed a difference between the two conditions, a point-to-point paired-t-test (significance level: 0.05) was applied to the HEPs across time with the LIMO toolbox (Pernet et al., 2011). This way, the time windows that showed a significant difference between both meditation sessions were detected in the time range from 0-800 ms. HEP amplitudes during these detected time windows at the Fz channel were then averaged for paired-t-tests (significance level 0.05, two-tailed) that compared the within-subject effect of meditation sessions. To test the topographic difference of HEPs between conditions in the detected time windows, a nonparametric randomization test using the Monte-Carlo method (10000 randomizations) was applied to estimate the significance probabilities for each channel without multiple corrections.

### 2.7. Control analysis of physiological recordings

To further exclude the possibility that the HEPs effect was caused by EEG artefacts related to heartbeats and breathing, a set of control analyses was completed including ECG analysis (average ECG amplitude in the HEPs time window, heart rate and heart rate variability) and respiratory analysis (respiration rate and respiration rate variability).

#### Surrogate heartbeats

To demonstrate that the HEP effects were synchronized with the heartbeats rather than resulting from random fluctuations of the amplitude, we generated 500 permutations of heartbeats separately for each condition following the previous studies (Babo-Rebelo et al., 2016; Park et al., 2016). To keep the same inter-beat interval and variability as the original R-peaks, the R-peaks were shuffled within each condition between trials, each trial lasting 20 seconds segmented from the 15-minute-long meditation session. The timing of the heartbeats in trial *i* was randomly assigned to trial *j*. After the permutations, the other analytical steps including the threshold of rejecting noisy trials remained the same as the aforementioned HEP analysis. Time windows that showed significance between two conditions were detected with the point-to-point paired-t-test (significance level: 0.05). The continuous time points that showed significance in the comparison between the 1PP and the 3PP condition were considered potential clusters. For each permutation, we extracted the largest positive sum of t values among the clusters, and compared the distribution of those surrogate values with the original sum of t values.

#### ECG analysis

After the R-peak detection described in the section on *ECG pre-processing*, the ECG signals were segmented into epochs of the same length for the HEPs analysis, i.e. [-100 800] ms. These epochs were averaged separately for 1PP and 3PP conditions, followed by the statistical test of average ECG amplitudes in the time window that showed the HEPs difference. Besides, the inter-beat intervals (IBI) were calculated as the time interval between two R-peaks. Time intervals smaller than 1500 ms or bigger than 500 ms (i.e. heartbeats at less than 40 or more than 120 bpm) were excluded. The number of heartbeats per minute during both meditation sessions was calculated as 60/(0.001*IBI). Additionally, to quantify the heart rate variability (HRV) (Shaffer & Ginsberg, 2017), the root mean square of successive heartbeats (RMSSD) was evaluated for each meditation condition.

#### Respiration analysis

The respiratory signals were first filtered with a high-pass filter of 0.2 Hz and a low-pass filter of 0.8 Hz, as usually done to exclude signals outside of normal respiration rates (12 ∼ 20 breaths per minute, Nicolò et al., 2020; Xu et al., 2021). Breathing peaks were detected by identifying the local maximum amplitudes of respiratory signals using the *BreathMetric* package (Noto et al., 2018). Similar to the ECG signals, the respiratory signals higher than a threshold of 10000 microvolts lasting longer than 200 s were excluded for further analysis. The breath rate variability was also measured with the root mean square of successive differences (RMSSD) (Kumar et al., 2020; Soni & Muniyandi, 2019).

### 2.8. Source estimation

To identify the brain areas where the HEP amplitudes differed across meditation conditions, the BrainStorm toolbox was used for the source estimation (Tadel et al., 2011) during the time window specified in the *HEPs statistical analysis* section. The head model was computed with OpenMEEG BEM based on the cortical surface of Colin27 MNI template, utilizing adaptive integration to enhance the precision (Gramfort et al., 2010). The sources of the HEP were determined using a weighted minimum-norm current estimate with the dSPM option where the dipole orientations were unconstrained for each meditation condition and each subject. Parameters included depth weighting with an order of 0.5 and a maximal amount of 10, a signal-to-noise ratio of 3, and a diagonal noise covariance. HEP differences were tested using a paired t-test with an alpha level of 0.01 without correction for multiple comparisons.

### 2.9. Statistics

For measures including the OBE-like sensations, 5D-ASC questions, PBBQ, and physiological measures, LME models were also applied to examine the effect of experimental conditions (1PP against 3PP). For the full-body illusion, we applied a Linear Mixed Effect (LME) model to examine the 2-by-2 design (experimental condition * visuo-tactile synchrony), meditation session and synchrony were set as fixed effects and the interaction between them was also explored. For all the abovementioned models, the participant number and the order of meditation sessions were set as random effects. To link the different measures including the full-body illusion, the PBB and the HEP, correlation analysis was conducted for the 3PP- 1PP effects while multiple comparisons were corrected with the Bonferroni approach.

## 3. Results

### 3.1. Stronger OBE-like sensations in the 3PP condition

Results of the OBE questionnaire (Wu et al., 2023) showed that participants experienced stronger OBE-like sensations in 3PP versus 1PP condition (Figure 3), as indexed by higher subjective ratings on the items of disconnection (*x*^2^(1) = 10.20, *p* = 0.0014, *BF*_10_ = 8.91), feeling of separation (*x*^2^(1) = 9.51, *p* = 0.002, *BF*_10_ = 7.19), and OBE-like feeling (*x*^2^(1) = 11.30, *p* = 0.00078, *BF*_10_ = 12.38). No other item of the OBE questionnaire showed significant differences between the two conditions (all p> 0.05).

**Figure 3.**
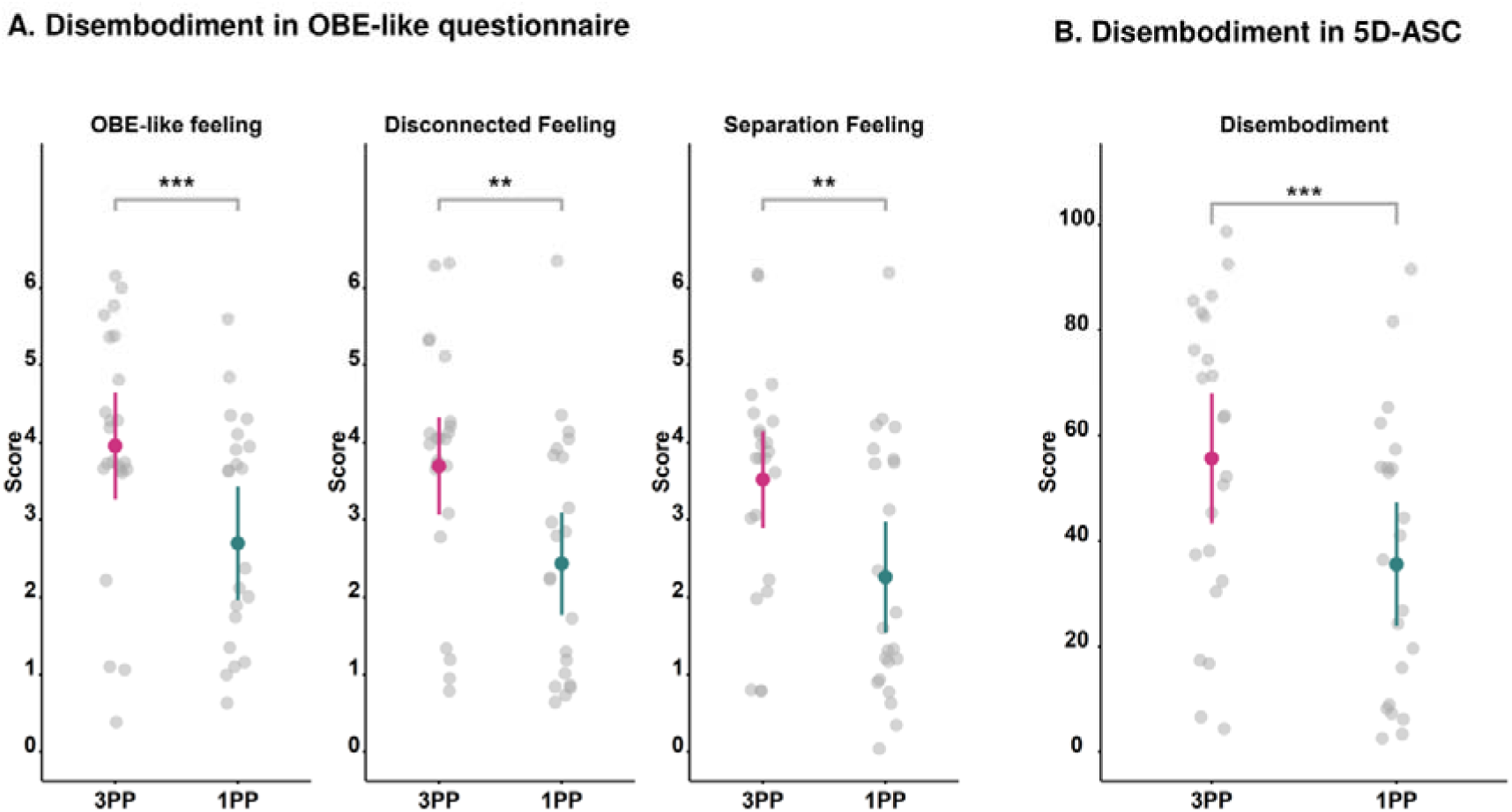
OBE-like sensations are stronger in the 3PP (pink) versus 1PP (green) condition. Data are shown for OBE-like feeling (A, left), disconnection (A, middle), separation (A, right) (range: 0-10) of the OBE-questionnaire (Wu et al., 2023) and the disembodiment item of the 5D-ASC questionnaire (Figure B, in percentage; range: 0-100).

Concerning the 5D-ASC, the dimension of disembodiment showed a significant difference (Figure 3B) between the 3PP condition (M = 55.70, SD = 28.57) and the 1PP condition (M = 35.58, SD = 27.05): *x*^2^(1) =11.33, *p* = 0.00076, *BF*_10_ = 12.55). The other ten dimensions did not show any difference across conditions (all *p* > 0.05; see Table 1).

**Table 1.** The subjective reported altered state of consciousness in percentage (%) according to the 5D-ASC (the dimension that showed significance (p<0.05) had been marked in bold font).

| No | Dimensions | 3PP condition | 1PP condition | LME |  |  | BF10 |
| --- | --- | --- | --- | --- | --- | --- | --- |
| | | M (SD) | M (SD) | df | $\chi^2$ | p | |
| 1 | Experience Of Unity | 40.72 (25.49) | 37.68 (22.01) | 22 | 1.17 | 0.28 | 0.36 |
| 2 | Spiritual Experience | 29.55 (23.65) | 25.01 (16.83) | 22 | 2.48 | 0.12 | 0.61 |
| 3 | Blissful State | 43.10 (30.08) | 37.39 (24.74) | 22 | 3.56 | 0.059 | 0.93 |
| 4 | Insightfulness | 38.22 (24.15) | 34.41 (22.67) | 22 | 1.51 | 0.22 | 0.42 |
| 5 | Disembodiment | 55.70 (28.57) | 35.58 (27.05) | **22** | **11.33** | **0.00076** | **12.55** |
| 6 | Anxiety | 19.86 (16.17) | 15.18 (14.31) | 22 | 1.97 | 0.16 | 0.50 |
| 7 | Complex Imagery | 31.81 (25.10) | 28.06 (25.95) | 22 | 1.31 | 0.25 | 0.38 |
| 8 | Elementary Imagery | 21.70 (23.96) | 17.49 (17.90) | 22 | 1.04 | 0.31 | 0.34 |
| 9 | Audio-Visual<br>Synthesiae | 30.19 (24.39) | 34.57 (25.38) | 22 | 1.57 | 0.21 | 0.43 |
| 10 | Changed Meanings of<br>Percepts | 37.52 (27.17) | 35.59 (22.59) | 22 | 0.27 | 0.60 | 0.25 |

### 3.2. Reduction of perceived body boundary in the 3PP condition

Changes in perceived body boundary were computed separately for each meditation condition as the difference of the PBBQ score measured before (baseline) and after each meditation condition. Both conditions showed a significant or marginal significant reduction in PBBQ (PBBQ score after meditations was lower than baseline, 3PP condition: *M* = −0.13, *SD* = 0.19, *t*(22) = −3.32, p = 0.003, *BF*_10_ = 12.13; 1PP condition: *M* = −0.07, *SD* = 0.16, *t*(22) = −2.04, *p* = 0.054, *BF*_10_ = 1.25). The reduction in the 3PP condition was significantly higher than in the 1PP condition (*x*^2^(1) = 6.46, *p* = 0.011). Bayes Factor however indicated that this result was inconclusive (*BF*_10_ = 2.65).

### 3.3. Reduced self-identification in the 3PP condition correlates with changes in perceived body boundary

In line with previous findings (Moon et al., 2024; Nakul et al., 2020; Lenggenhager et al., 2007), we replicated the main effect of stroking synchrony (*x*^2^(1) = 33.04, *p* = 9.05* 10^-9^) on self-identification. Moreover, the 3PP condition significantly reduced self-identification compared to the 1PP condition (main effect of experimental condition: *x*^2^(1) = 11.68, *p* = 6.32 * 10^-4^) and there was also an interaction between experimental condition and visuo-tactile stroking condition (*x*^2^(1) = 4.08, *p* = 0.043). Post-hoc testing indicated significantly lower self-identification during the 3PP vs. 1PP condition during asynchronous stroking (3PP condition: *M* = 4.52, *SD* = 1.79; 1PP condition: *M* = 5.28, *SD* = 1.64; *t*(45) = 3.24, *p* = 0.0023, *BF*_10_ = 14.11), which was absent during synchronous stroking (3PP condition: *M* = 5.61, *SD* = 1.56; 1PP condition: *M* = 5.80, *SD* = 1.29; *t*(45) = 1.39, *p* = 0.17, *BF*_10_ = 0.39). Thus, self-identification with the seen avatar was weaker in the 3PP condition and this was particularly the case during asynchronous stroking.

Concerning mislocalization of touch there was a main effect of synchrony (*x*^2^(1) = 203.78, *p* = 2.0 * 10^-16^) with significantly larger ratings in the synchronous vs. asynchronous visuo-tactile condition, as reported previously (i.e., Lenggenhager, Tadi, et al., 2007). There was no main effect of intervention (*x*^2^(1) = 0.57, *p* = 0.45) nor interaction effect (*x*^2^(1) = 1.68, *p* = 0.19). For the control question, there were no main effects (visuo-tactile synchrony: *x*^2^(1) = 3.41, *p* = 0.065; experimental condition: *x*^2^(1) = 7.7 * 10^-3^, *p* = 0.93) and no interaction (*x*^2^(1) = 0.63, *p* = 0.43).

Additionally, we explored whether the changes in self-identification (during the full-body illusion task) correlated with the PBBQ scores. For this, we compared the difference in self-identification with the PBBQ scores between conditions. Correlation analysis revealed a positive correlation between the PBBQ difference between conditions and self-identification for synchronous stroking condition of the full-body illusion (*r* = 0.52, *p* = 0.01, see Figure 5). Thus, the stronger participants identified with the seen body (self-identification), the stronger they reported a transparent body boundary in the 3PP versus 1PP condition. This correlation was absent for asynchronous stroking (*r* = −0.15, *p* = 0.48).

**Figure 4.**
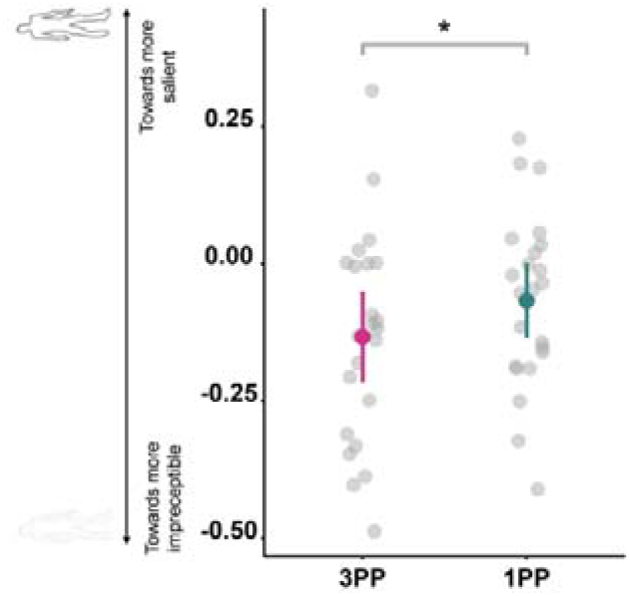
Difference scores of the perceived body boundary after both meditation sessions compared to the baseline (zero score represents a score equal to baseline; a negative value a more blurred body boundary). Data show a greater reduction of the perceived body boundary in the 3PP condition (pink) compared to the 1PP condition (green).

**Figure 5.**
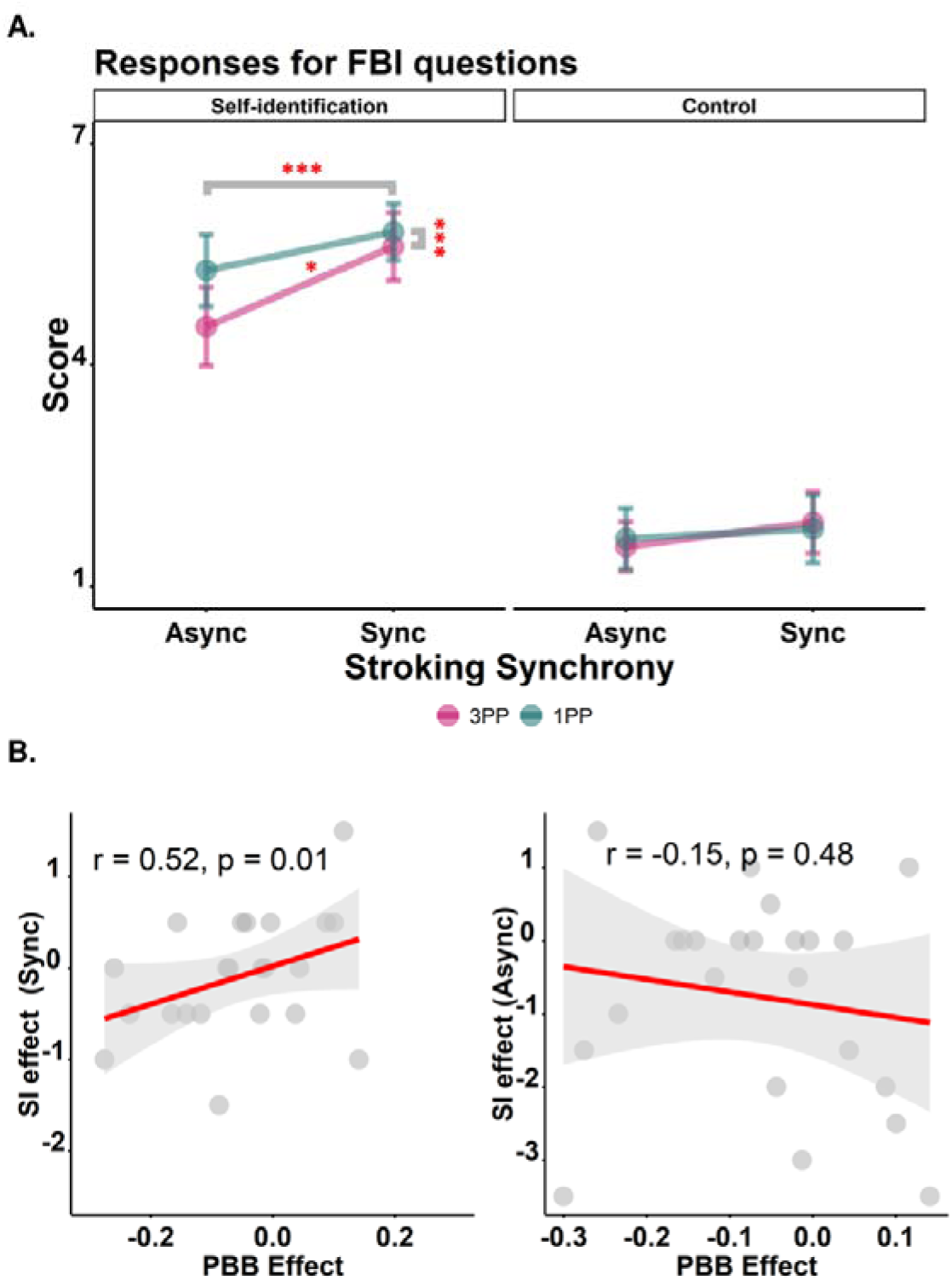
A. Results for self-identification (left) and a control question (right) are shown for synchronous and asynchronous stroking for the 3PP (pink) and 1PP condition (green). Dots represent the average scores and the error bars are the 95% confidence interval. The significance of main effects and interaction is indicated (‘*’ represents p<0.05, ‘***’ represents p<0.001). B. Difference in self-identification ratings (3PP - 1PP) correlates with the difference in perceived body boundary (PBBQ difference between conditions). This was found in the synchronous stroking condition (left), but not for the asynchronous stroking condition.

### 3.4. Stronger, more negative, HEPs in the 3PP condition

First, EEG analysis revealed a prominent HEP at electrode Fz (Figure 6A), similar to previous HEP work (Fukushima et al., 2011; Park et al., 2016; Tanaka et al., 2023). Second, the averaged HEP waveform at electrode Fz in the 3PP condition was characterized by a significantly more negative amplitude in a time window ranging from 313 to 754 ms (timepoint-by-timepoint t-test from 0-800 ms after the heart beat; p<0.05) (Figure 6A). Third, full-scalp analysis during this time window revealed two fontal electrodes (Fz, FCz) differing significantly between both conditions (p<0.01) (permutation t-test; Monte-Carlo). The average amplitudes at both electrodes for this time period were further tested with a paired-t-test, showing a more negative HEP amplitude in the 3PP condition (Fz: M = −0.21, SD = 1.70; FCz: M = −0.21, SD = 1.70) compared to the 1PP condition (Fz: M = 0.56, SD = 1.58; FCz: M = 0.56, SD = 1.58) at electrode Fz (*x*^2^(1) = 13.28, *p* = 0.00027, *BF*_10_ = 21.89) and FCz (*x*^2^(1) = 9.41, *p* = 0.0022, *BF*_10_ = 21.88).

**Figure 6.**
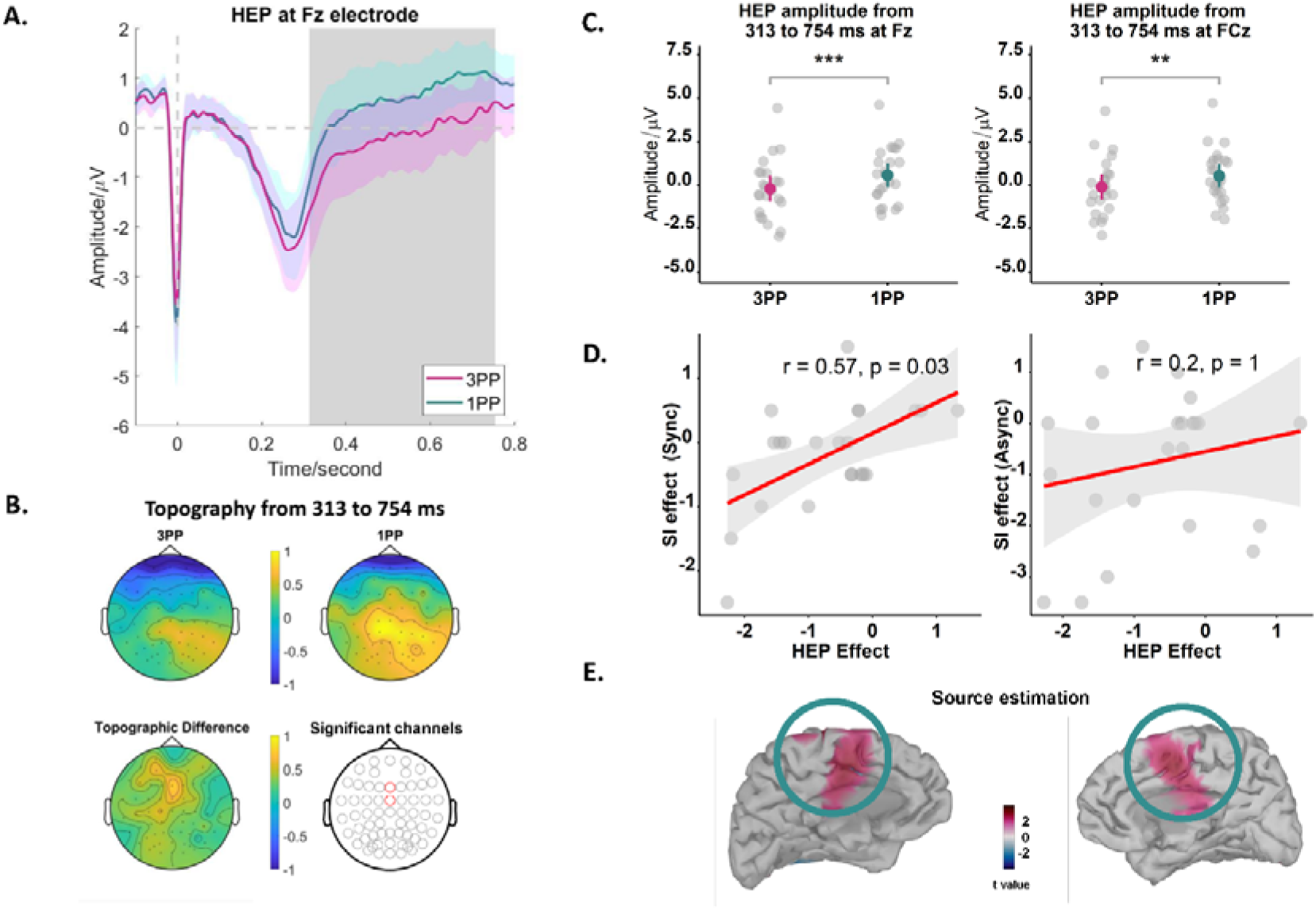
A. HEPs are significantly lower amplitude in the time window of 313 – 754 ms (marked grey) in the 3PP vs. 1PP condition. The pink and green shadowed areas represent the 95% confidence interval separately for 3PP and 1PP condition. B. Topography of HEP in the 3PP condition (left), the 1PP condition (middle), and for the difference between both conditions (right). C. Boxplot of HEP amplitude at Fz electrode for both conditions. Significant difference was found only in two frontal sites (Fz, FCz). (D) The difference in self-identification ratings (3PP - 1PP) correlates with the difference in HEP amplitude (HEP difference between conditions). This correlation was found in the synchronous stroking condition (left), but not for the asynchronous stroking condition. (E) Source estimation for the HEP from 313 to 754 ms.

Additionally, we explored whether changes in self-identification during the full-body illusion were associated with changes in HEP amplitude during meditation. Specifically, we compared the differences in self-identification with the HEP amplitude differences between conditions (average amplitude of Fz and FCz electrode). Correlation analysis indicated a significant positive relationship between the HEP difference and self-identification in the synchronous stroking condition (*r* = 0.57, *p* = 0.03; see Figure 6D). This finding suggests that a stronger identification with the seen body (self-identification) corresponds to enhanced neural responses in the 3PP versus 1PP condition. In contrast, no significant correlation was observed for the asynchronous stroking condition (*r* = 0.2, *p* = 1) after correcting for multiple comparisons with Bonferroni in which the p values were multipled by the number of comparisons (Curtin & Schulz, 1998).

Finally, to identify the brain regions reflecting the significant condition differences in HEP amplitude at frontocentral scalp electrodes (Figure 6B), we reconstructed the sources of HEP during the time window from 313 to 754 ms separately for the 3PP condition and 1PP condition. This analysis localized HEP differences in the posterior cingulate cortex (paired t-test; p < 0.01) (PCC, see Figure 6E) and the medial prefrontal cortex (mPFC).

### 3.5. HEP Control analysis

#### Surrogate R-peaks results

The summed t values of the clusters were computed for the 500 permutations and the largest cluster was picked by selecting the cluster with the largest summed t value at each permutation. None of the 500 permutations showed a summed t value as large as the original ones obtained from the HEP time-locked to real R-peaks (Figure 7A), confirming that the differential HEP effect is locked to the heartbeat and not due to random fluctuations of the HEP amplitude.

**Figure 7.**
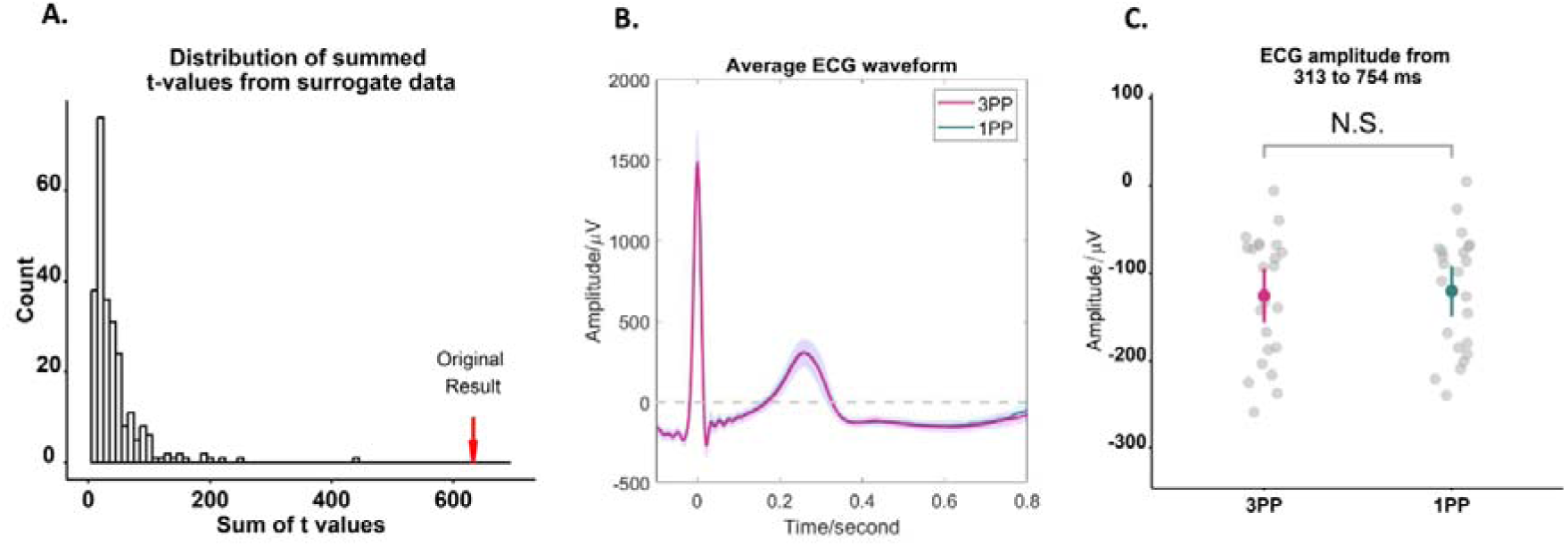
A) The histogram of the distribution of the maximal summed t values, derived from 100 permutations of surrogate heartbeats. Notably, the original summed t value (indicated by the arrow) lies outside the distribution of statistics generated from surrogate data. B) The average ECG waveforms in the HEPs time window (−100 to 800 ms) during the 3PP and 1PP conditions. C) Mean and confidence interval of the ECG amplitudes in the specified time window (313 to 754 ms).

#### ECG results

Further analysis revealed that these HEP changes cannot be accounted for by differences in ECG parameters. Control analysis, conducted to examine potential cardiac confounds (ECG waveforms were obtained by averaging the ECG amplitudes, Figure 7B), showed no differences between the 3PP and the 1PP condition in the ECG amplitude in the same time window that showed a HEP difference, i.e., ranging from 313 to 754 ms (Figure 7C, see Table 2 for the detailed descriptive and statistical tests).

**Table 2.**
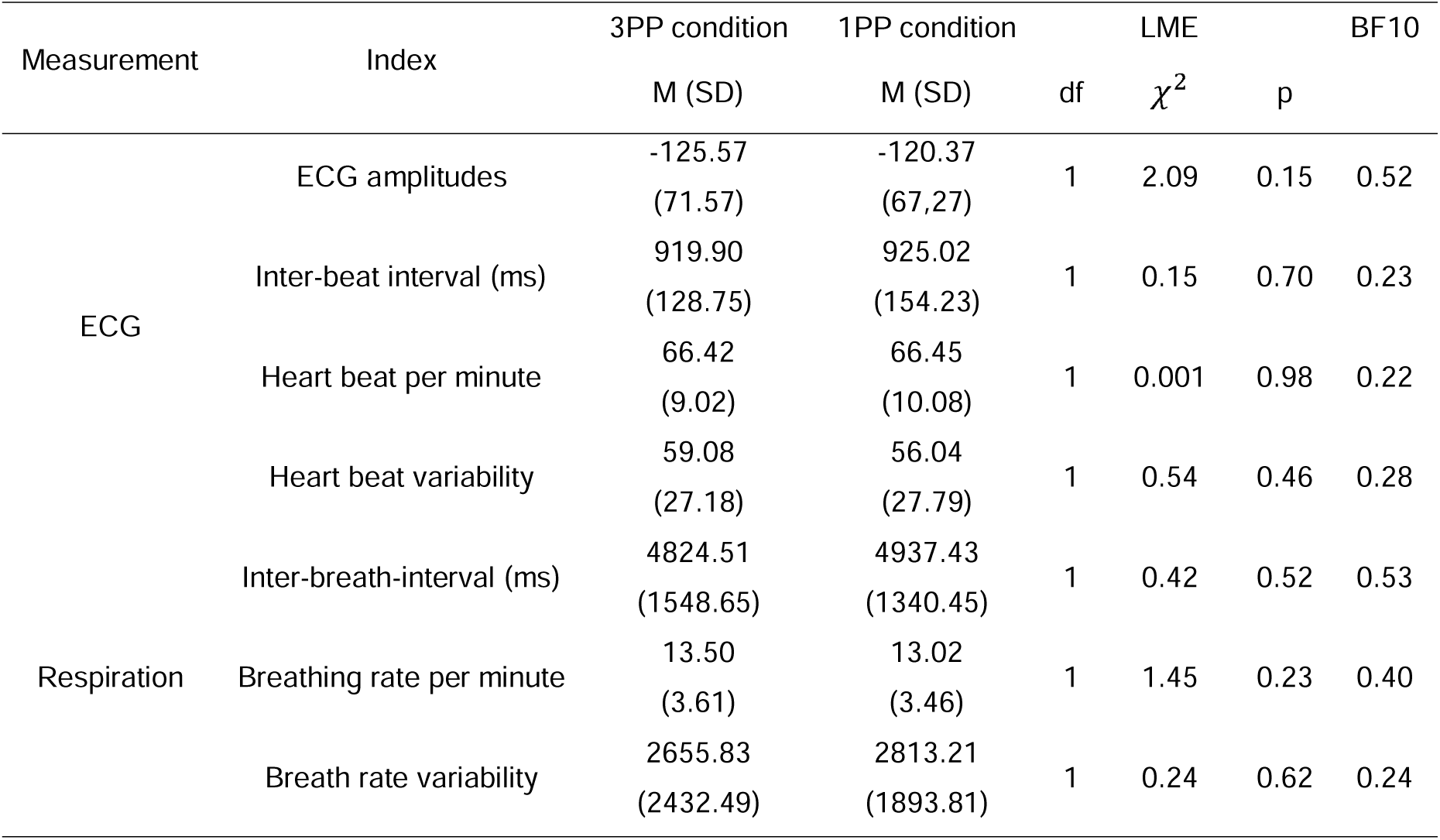
Descriptive and statistical results of the control analysis of the HEP, including ECG and respiratory signals.

Beyond the ECG amplitudes, we measured heart rate and heart rate variability during two sessions to exclude the confound of cardiac changes. Both the inter-beat interval and the heart rate did not show a difference between meditation sessions. The heart rate variability was examined with the root mean square of successive R-R interval (RMSSD). Results showed that the 3PP condition showed no difference with the 1PP condition on the heart rate variability (see Table 2).

#### Respiratory results

The breathing signals were extracted for the 3PP condition and 1PP condition separately. The breathing rate per minute did not show a difference between the 3PP and the 1PP condition. The pattern remained the same when calculating the inter-breath-interval. Also, the breath rate variability was compared against meditation sessions and no effect had been found, by means of RMSSD.

## 4. Discussion

Extending previous research on the minimal self in meditation practice (Canby et al., 2024; Dor-Ziderman et al., 2016; Droit-Volet & Dambrun, 2019; Lindahl & Britton, 2019), we here leveraged a new VR platform to perform guided meditation while experimentally manipulating the minimal self (3PP versus 1PP) and obtaining behavioral minimal self measures (changes in self-identification) and EEG measures. We report that the 3PP versus 1PP condition was associated (1) with stronger feelings of separation, detachment, and OBE-like sensations (5D-ASC, OBE questionnaire) and weaker self-identification with the seen body (full-body-illusion). The 3PP condition also (2) led to a more transparent body boundary (PBBQ), as reported previously for meditation practice and the magnitude of PBBQ changes between conditions positively correlated with the self-identification changes in the full-body-illusion-inducing synchronous condition. (3) HEP data revealed differential neural responses between both conditions, characterized by a more negative HEP amplitude in the 3PP condition(Park et al., 2016, 2018), linked to activation in PCC and mPFC, two structures that have previously been linked to HEPs and the minimal self. (4) Control analyses show that these changes were not caused by physiological changes including heart rate, heart beat variability, breath rate, and breath rate variability.

### 4.1. VR-supported meditation practice induces stronger disembodiment and reduced salience of the perceived body boundary

To manipulate the minimal self during meditation, we combined meditation practice with a VR platform that immersed participants in a virtual environment where they could view a 3D representation of their own bodies. The platform enabled us to employ different immersive perspectives in each condition. In the 3PP condition, we combined a visual transition from a first-person to third-person viewpoint with open monitoring meditation during which participants were instructed to take a de-centered and distanced perspective and witness moment-to-moment fluctuations of conscious experience. In the 1PP condition, participants were exposed to a transition of the shelter area within the virtual forest (same speed-distance as 3PP transition), while remaining in a first-person viewpoint and instructed to engage in focusing on the environment (forest). Data show that 3PP versus 1PP condition was associated with stronger feelings of separation/detachment of our participants’ self from their body and with other OBE-like sensations (OBE questionnaire; Wu et al., 2024; disembodiment items from 5D-ASC). Thus, integrating a 1PP to 3PP shift in perspective into meditation was associated with OBE-like sensations during meditation practice, comparable to such sensations reported in different experimental setups manipulating visuo-tactile in healthy individuals (Ehrsson, 2007; Ionta et al., 2011; Pfeiffer et al., 2013, 2016) or visuo-vestibular signals (Wu et al., 2024). They also correspond to disembodiment scores in the 5D-ASC (Schmidt & Berkemeyer, 2018), though less intense than reported after intake of psychedelic substances (i.e., lysergic acid diethylamide; Kraehenmann et al., 2017). The present OBE-sensations were also weaker than those reported in OBEs of neurological patients (Blanke et al., 2002, 2004, 2005; Hiromitsu et al., 2020) or spontaneous OBEs in healthy participants (Braithwaite et al., 2011; Milne et al., 2019).

Moreover, the 3PP versus 1PP condition was accompanied by a less salient body boundary (PBBQ), as reported previously for meditation practices such as body scan meditation (Dambrun, 2016; Dambrun et al., 2019) or mindfulness training (Ataria et al., 2015), in novices (Hanley et al., 2020) and experts (Linares Gutiérrez et al., 2022). These data show that the 3PP condition induced a decreased, more transparent body boundary that has been argued to be of relevance for the minimal self in meditation practices (Ataria et al., 2015; Dambrun, 2016; Lindström et al., 2023; Nave et al., 2021, 2021; Trautwein et al., 2024) or intake of psychedelic substances (Canby et al., 2024). A large body of meditation research suggests that maintaining a distanced perspective on one’s experiences is a crucial aspect of de-centering (i.e., Bennett et al., 2021; Kessel et al., 2016). A novel aspect of the present VR-enabled 3PP condition is that such distancing during open monitoring can be additionally supported by ‘visual distancing’ of the practicioner’s viewpoint, potentially enhancing the ability to de-center and to re-perceive events in conscious experience, a metacognitive ability commonly cultivated by mindfulness meditators (Dahl et al., 2015; Gecht et al., 2014; Hölzel et al., 2011; Kessel et al., 2016). There is a wide variate of mindfulness practices (Dunne, 2011) and these likely tap into different behavioral and neural mechanisms (Lutz et al., 2015). Our VR-based approach may help in investigating differences between such contenmplative traditions and study common and shared mechanisms of mental and visual distancing in meditation practice as well as their potential to jointly support de-centering in meditation practice.

### 4.2. Behavioral changes of the minimal self in the full-body illusion

We also observed condition-related changes in the full-body illusion. Following both meditation conditions, our participants were sensitive to visuo-tactile stimulation, showing similar effects as observed previously (higher mislocation of touch; higher self-identification in synchronous-vs.-asynchronous condition) (Aspell et al., 2009; Bayer et al., 2023; Lenggenhager et al., 2007; Noel et al., 2015; Park et al., 2016). Critically, self-identification was differently modulated by our meditation conditions and characterized by lower self-identification in 3PP vs. 1PP condition. Moreover, this differential decrease across conditions was driven by lower values during the asynchronous 3PP condition that were equally strong between 3PP and 1PP during synchronous stimulation (Figure 5), suggesting that 3PP meditation disrupts multisensory processes of the full-body illusion, especially during asynchronous stroking that are overcome by heightened sensitivity during synchronous stroking. As reported previously for meditation practice, the present 3PP condition also led to a more transparent body boundary and the magnitude of these changes between conditions positively correlated with the self-identification changes in the illusion-inducing synchronous condition. These findings further associate the perceived transparency of the body boundary to a classical measure of multisensory own body perception (full-body illusion) and more generally link the minimal self, as studied in cognitive neuroscience, to meditation practice, extending previous data using questionnaires and detailed verbal interviews (Berkovich-Ohana et al., 2020; Nave et al., 2021).

### 4.3. More negative heartbeat evoked response in 3PP meditation

Neural responses time-locked to heartbeats, HEPs, have been shown to reflect changes in self-identification induced by visuo-tactile stimulation (Park et al., 2016). Here we report that both conditions showed prominent HEPs characterized by typical topography and that the HEP in the 3PP versus 1PP meditation condition (associated with decreased self-identification) showed a more negative HEP. HEP decrease has been observed in the full-body illusion, during asynchronous visuo-tactile stimulation and was associated with decreased self-identification as the present 3PP condition (Park et al., 2016). This previous work, however, did not investigate meditation practice. HEP changes have further been associated with visual consciousness(Park & Tallon-Baudry, 2014), narrative self (Babo-Rebelo et al., 2016), and long-term meditation practice (i.e. greater HEP amplitudes in Tibetan Buddhist meditators versus controls) (Jiang et al., 2020). Methodologically, we suggest that HEPs are an interesting neuromarker for meditation research, because it can be acquired during meditation practice without exposing practicioners to additional sensory stimulation or cognitive load, often required by other evoked potentials(i.e., Chan et al., 2020; Singh & Telles, 2015).

Concerning HEP topography and its neural sources, our data highlighted fronto-central regions showing 1PP-3PP differences, aligning with meta-analytical HEP data (Coll et al., 2021) that arguably reflect the involvement of PCC and interoceptive cortices (Al et al., 2021; Gentsch et al., 2019; Hodossy et al., 2021; Kumral et al., 2022). The PCC has been associated with self-referential processing (Agathos et al., 2023; Brewer et al., 2013; Leech & Sharp, 2014; Parvizi et al., 2021) and the minimal self (Lyu et al., 2023; Park et al., 2016; Vesuna et al., 2020). The PCC is also commonly activated across various meditation practices (Brewer & Garrison, 2014; Kral et al., 2019, 2022; Zsadanyi et al., 2021), and has been consistently linked to the HEP (Jiang et al., 2020). Furthermore, the PCC has been recently associated with the experience of ‘boundarylessness’ (Dor-Ziderman et al., 2016; Lindström et al., 2023), a meditation experience that parallels the perceived body boundary changes reported in our study. Additional activation was observed in the mPFC in the 3PP versus 1PP condition, consistent with the role of this region in self-referential processing (i.e., Qin et al., 2020).

To summarize, the present 3PP practice – meditating while perceiving the world and one’s body from a distanced backward shifted third-person viewpoint — induces disembodiment, a less clearly experienced body boundary, and decreased self-identification with one’s body that depends on the synchrony of visuo-tactile stimulation. The link between self-identification and perceived body boundary following 3PP meditation was corroborated by our additional finding that the PBBQ decrease between conditions positively correlated with the self-identification changes in the full-body-illusion-inducing synchronous condition, linking the perceived salience of the body boundary to a measure of the minimal self. These subjective-behavioral changes were also reflected in HEP-related differential activation of PCC and mPFC. Using a VR-supported meditation platform and methodologies, our study reveals novel connections between the sense of self in meditation practice and the neuroscience of the multisensory bodily self, reflected in subjective and behavioral changes in multisensory own body perception, and associated interoceptive bodily processes.

## Funding

This research is supported by All Here SA (The Home Within: Minimal Phenomenal Selfhood, Phenomenal Existence, and Meditation Neuroengineering). OB is supported by Carigest SA, Swiss National Science Foundation (3100A0-112493), Bertarelli Foundation.

## CRediT authorship contribution statement

(HY - Hang Yang, BH - Bruno Herbelin, CN - Chuong Ngo, LV - Loup Vuarnesson, OB - Olaf Blanke)

HY: Conceptualization, Data curation, Formal Analysis, Investigation, Methodology, Software, Validation, Visualization, Writing– original draft, Writing – review & editing.

BH: Conceptualization, Data curation, Methodology, Project administration, Resources, Visualization, Supervision, Writing – review & editing.

CN: Methodology, Investigation.

LV: Methodology, Software, Investigation, Visualization.

OB: Conceptualization, Funding acquisition, Methodology, Project administration, Resources, Supervision, Validation, Writing– original draft, Writing – review & editing.

## Acknowledgments

We thank Erkin Bek for guidance and discussion from the beginning of the project and Fosco Bernasconi and Mariana Babo-Rebelo for their advice in data analysis. We thank the Foundation of Campus Biotech Geneva (FCBG) for providing help and support with the EEG system. We thank all participants for their participation.

## Notes

### Competing Interest Statement

The authors have declared no competing interest.

